# HuR -dependent SOD2 protein synthesis is an early adaptation of ovarian cancer cells in response to anchorage independence

**DOI:** 10.1101/2021.02.19.432010

**Authors:** Yeon Soo Kim, Jaclyn E. Welles, Priscilla W. Tang, Zaineb Javed, Amal T. Elhaw, Karthikeyan Mythreye, Scot R. Kimball, Nadine Hempel

**Author notes:** **Corresponding Author**: Nadine Hempel, Ph.D., Department of Pharmacology, Penn State University College of Medicine, MC R130, 500 University Drive, Hershey PA 17033-0850.

## Abstract

The presence of malignant ascites is a common feature of advanced stage ovarian cancer. During metastasis, as cells detach from the tumor and extravasate into the peritoneal fluid, ovarian cancer cells must adapt to survive the loss of anchorage support and evade anoikis. An important pro-survival adaptation in this context is the ability of tumor cells to increase their antioxidant capacity and restore cellular redox balance. We previously showed that the mitochondrial superoxide dismutase SOD2 is necessary for ovarian cancer cell anoikis resistance, anchorage-independent survival and spheroid formation, and intraperitoneal spread *in vivo*. We now demonstrate that the upregulation of SOD2 protein expression is an early event initiated in response to anchorage independence and occurs at the post-transcriptional level. SOD2 protein synthesis is rapidly induced in the cytosol within 2 hours of matrix detachment. Polyribosome profiling demonstrates an increase in the number of ribosomes bound to SOD2 mRNA, indicating an increase in *SOD2* translation in response to anchorage-independence. Mechanistically, we find that anchorage-independence induces cytosolic accumulation of the RNA binding protein HuR/ELAVL1, leads to HuR binding to *SOD2* mRNA, and that the presence of HuR is necessary for the increase in *SOD2* mRNA association with the heavy polyribosome fraction and SOD2 protein synthesis. Cellular detachment activates the stress-response protein kinase p38 MAPK, which is necessary for HuR-*SOD2* mRNA binding and the rapid increase in SOD2 protein expression. Moreover, HuR is necessary for optimal cell survival during early stages of anchorage independence. These findings uncover a novel post-transcriptional stress response mechanism by which tumor cells are able to rapidly increase their mitochondrial antioxidant capacity to adapt to stress associated with anchorage-independence.

## Introduction

Despite significant improvements in our understanding of the genomic and RNA expression signatures of ovarian cancer, the 5-year survival rate of patients has remained relatively low (Torre *et al*, 2018). Epithelial ovarian cancer (EOC), which accounts for 90% of ovarian cancer cases, remains the most deadly gynecological malignancy, partly due to the asymptomatic progression to advanced disease. About 80% of patients with high grade serous adenocarcinoma, the most common subtype of EOC, are diagnosed at stage III and stage IV. At this stage the 5-year survival rate remains less than 30% (Torre *et al*., 2018), and is characterized by significant tumor spread throughout the peritoneal cavity and the accumulation of malignant ascites (Ahmed & Stenvers, 2013). This peritoneal fluid is thought to further facilitate transcoelomic metastasis of ovarian cancer by providing a medium in which extravasating cells disseminate throughout the peritoneal cavity where they eventually invade the mesothelial wall of the peritoneum, and neighboring organs including the omentum, liver, intestines, and pleural fluid.

Disseminated ovarian tumor cells undergo stress adaptation to survive in non-adherent conditions. One adaptation to evading anchorage-independent cell death, known as anoikis, is by increased antioxidant capacity to reduce the detachment-induced surges in reactive oxygen species (ROS) (Jiang *et al*, 2016; Schafer *et al*, 2009). *In vivo* studies have demonstrated that increased antioxidant enzyme expression and small molecule antioxidant treatment promotes the metastatic spread of melanoma and breast cancer cells (Davison *et al*, 2013; Piskounova *et al*, 2015), suggesting that the maintenance of redox homeostasis is a key adaptation during metastasis. Similarly, we have shown that ovarian cancer cells increase their mitochondrial antioxidant capacity after matrix detachment, by upregulating the expression and activity of the deacetylase sirtuin 3 (SIRT3), and its target protein the mitochondrial superoxide dismutase SOD2 (Kim *et al*, 2020). Our previous work demonstrated that SIRT3-dependent SOD2 deacetylation is necessary for SOD2 activity and that upregulation of both SIRT3 and SOD2 are necessary for anoikis resistance and *in vivo* transcoelomic spread of ovarian cancer cells (Kim *et al*., 2020).

SOD2 is a nuclear encoded mitochondrial protein that is responsive to stress-activated transcriptional regulation (Kim *et al*, 2017). An extensively studied stress-response transcription factor implicated in the regulation of antioxidant responses of tumor cells is Nrf2/NFE2L2, which has been implicated with the increases in SOD2 expression observed in breast cancer and clear cell ovarian carcinomas (Hart *et al*, 2016; Hemachandra *et al*, 2015; Konstantinopoulos *et al*, 2011). *SOD2* transcription is also induced by the sirtuin regulated transcription factor Foxo3A (Kenny *et al*, 2017), and by NF-κB, which has also been implicated in the upregulation of *SOD2* transcription in response to matrix detachment of breast cancer cells (Kamarajugadda *et al*, 2013). Although much emphasis has been placed on the SIRT3-dependent regulation of SOD2 activity and stress response pathways leading to *SOD2* transcription in cancer, whether translational regulation contributes to SOD2 expression in tumor cells has not been investigated in depth.

Posttranscriptional and translational regulatory mechanisms are crucial for fine-tuning of gene expression. In particular, the interplay between mRNAs, miRNAs, and RNA-binding proteins has been implicated in cancer development and metastasis (Audic & Hartley, 2004; van Kouwenhove *et al*, 2011; Wurth & Gebauer, 2015). Among RNA binding proteins, HuR exerts many effects such as increased mRNA stability and translational efficiency by binding to the AU-and U-rich elements (AREs) in the 3’ UTR of target mRNAs (Abdelmohsen & Gorospe, 2010). Following activation HuR translocates from the nucleus to the cytoplasm, where it binds and regulates mRNAs that encode proteins involved in oncogenic signaling pathways (Epis *et al*, 2011; Mazan-Mamczarz *et al*, 2008), anti-apoptosis (Filippova *et al*, 2011), cell cycle (Lal *et al*, 2014; Wang *et al*, 2000b), and chemoresistance (Raspaglio *et al*, 2010), all of which converge on accelerated tumorigenesis and aggressive phenotypes (Wang *et al*, 2013). Importantly, HuR can translocate to the cytoplasm upon genotoxic or environmental stimuli in cancer cells (Lafarga *et al*, 2009; Lal *et al*., 2014), which suggests that HuR-dependent translation may be a critical stress adaptation utilized by cancer cells. HuR expression analyses across different malignancies including ovarian cancer showed that its cytoplasmic accumulation is correlated with advanced stages and poor prognosis (Denkert *et al*, 2004a; Denkert *et al*, 2004b; Miyata *et al*, 2013; Mrena *et al*, 2005). However, it is not known if HuR is involved in ovarian cancer metastatic spread, or specifically upregulated as an adaptation to anchorage independent stress.

Interestingly, a transcriptome-wide RNA-binding analysis identified multiple HuR binding sites in the 3’ UTR of SOD2 mRNA (Lebedeva *et al*, 2011), but the functional consequences of these sites related to SOD2 mRNA translation remain elusive. Given that HuR is a stress responsive RNA binding protein promoting cell survival, we investigated if SOD2 mRNA is a target of HuR in tumor cells. In the present work, we show for the first time that SOD2 mRNA is a target of HuR in ovarian cancer and that the interaction of HuR with *SOD2* mRNA is required for rapid *de novo* SOD2 protein synthesis after matrix detachment in a p38 MAPK dependent manner. Our study provides evidence for a novel mechanism of SOD2 regulation in response to acute stress associated with anchorage-independence. It demonstrates that tumor cells are able to increase their antioxidant defenses at multiple levels beyond transcription, allowing for rapid adaptations to stress associated with different stages of tumor metastasis.

## Results

### SOD2 protein expression increases rapidly in response to anchorage independence

In previous work, we demonstrated that SOD2 activity increases in ovarian cancer cells within 2 hours in ultra-low attachment (ULA) cell culture conditions, which is followed by increased transcription within 24 hours in anchorage-independence (Kim *et al*., 2020). In addition to the rapid increase in SOD2 superoxide dismutase activity, which we reported to be dependent on SIRT3 (Kim *et al*., 2020), it was noted that the cytosolic SOD2 protein pool rapidly increases following detachment (Fig 1A). Subcellular fractionation demonstrated an average 4.7-fold increase in OVCA433 cytosolic SOD2 expression after 0.5 hour of cell detachment compared to attached cells, while a 1.5-fold increase was observed after 2 hours in anchorage-independent conditions in OVCAR10 cells (Fig 1A&B). In OVCA433 anchorage-independent cultures *SOD2* mRNA increases trailed the changes in SOD2 protein expression, suggesting that the rapid rise in SOD2 protein following detachment is likely independent of increases in transcription in this cell line (Fig 1C, Supp Fig 3B). To determine if the observed increase in cytosolic SOD2 represents a newly synthesized SOD2 protein pool, cells were treated with the protein synthesis inhibitor cycloheximide, which decreased total SOD2 protein levels under anchorage independence (Fig 1D). ^35^S-Met/Cys incorporation assays demonstrated a global increase in protein synthesis immediately following detachment (Suppl. Fig 1A), and subsequent immunoprecipitation of SOD2 confirmed enhanced synthesis of SOD2 protein in anchorage independence. After 2 hours of incubation, ovarian cancer cells demonstrated 1.8-fold (OVCA433) and 2.4-fold (OVCAR10) increases in ^35^S-Met/Cys incorporation into the SOD2 protein compared to attached conditions, which was abrogated by cycloheximide.

**Figure 1.**
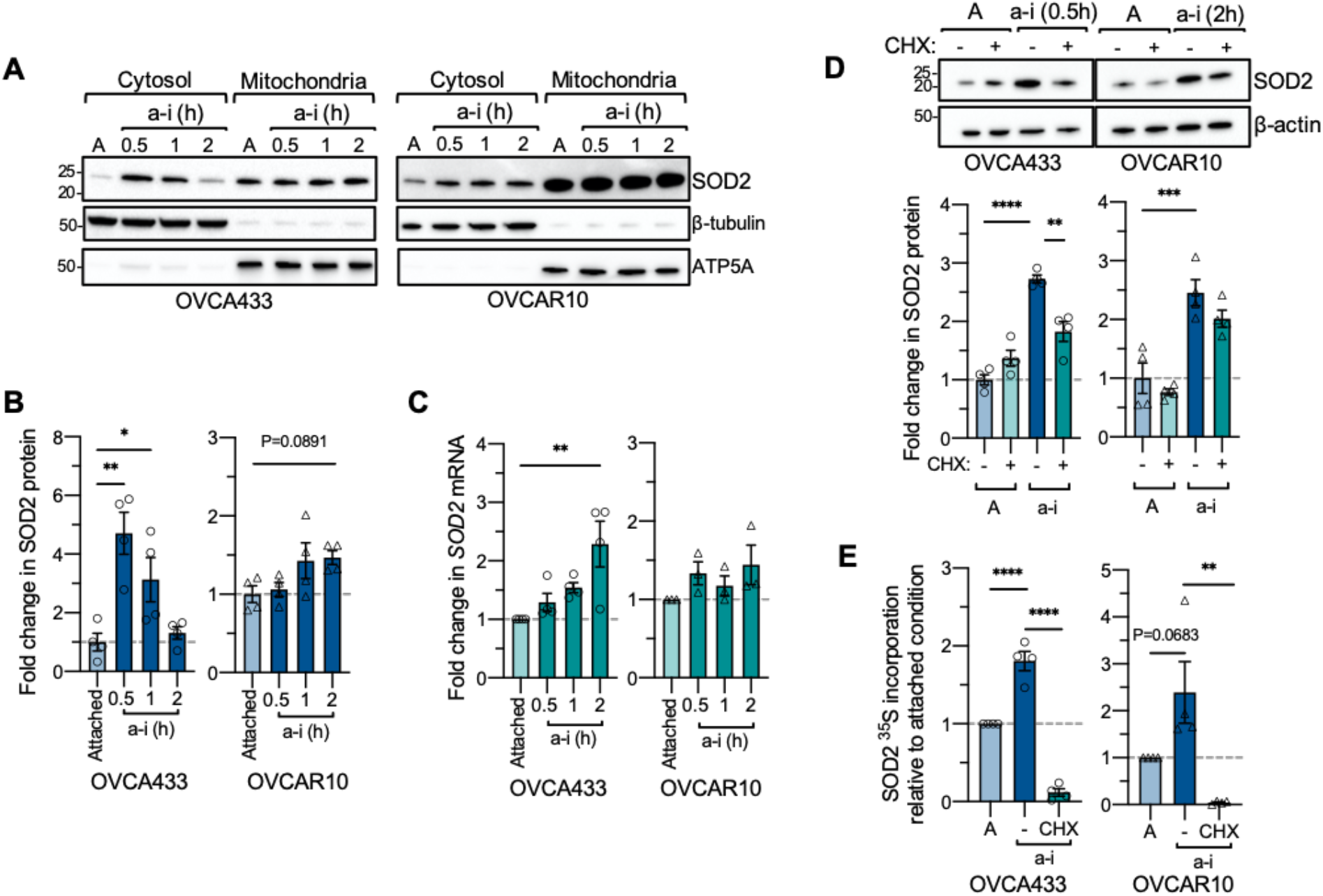
A. The cytosolic SOD2 protein pool increases rapidly in response to anchorage-independence (a-i), compared to attached culture conditions (A). Cells were maintained for indicated times in ULA plates and SOD2 protein expression assessed following cellular fractionation and immunoblotting. B. Fold change in SOD2 cytosolic protein expression in response to anchorage-independent (a-i) culture was quantified using densitometry, normalized to β-tubulin loading control and expressed relative to attached (A) culture conditions (*n*=4, one-way ANOVA, OVCA433 *P*=0.0015, OVCAR10 *P*=0.0744, Dunnett’s multiple comparison test \**P*<0.05; \*\**P*<0.01). C. Fold change in *SOD2* mRNA in response to short term anchorage-independent culture was assessed using semi-quantitative real time RT-PCR (*n*=3-4, one-way ANOVA, OVCA433 *P*=0.0069, OVCAR10 *P*=0.2946, Dunnett’s multiple comparison test \**P*<0.05; \*\**P*<0.01). D. Total SOD2 protein levels were assessed by immunoblotting in response to culture in anchorage-independent conditions and protein synthesis inhibited by cycloheximide (CHX, 20 µg/mL; *n*=4, one-way ANOVA, *P*<0.0001, Tukey’s multiple comparison test \**P*<0.05; **P<0.01). E^35^S-Met/Cys incorporation assay followed by SOD2 IP (Suppl. Fig 1B&C), demonstrates increased ^35^S-Met/Cys incorporation into SOD2 under anchorage independence compared to attached cells, which is abrogated in the presence of cycloheximide (*n*=4, one-way ANOVA, OVCA433 *P*<0.0001, OVCAR10 *P*=0.0057, Tukey’s multiple comparison test \*\**P*<0.01; \*\*\*\**P*<0.0001).

To confirm that the increases in SOD2 protein expression are due to *de novo* protein synthesis, ribosome-mediated mRNA translation was assessed using polyribosome profiling. Following centrifugation, sucrose gradients were separated into four fractions and RNA was isolated from each fraction. Fraction 1 contained mRNAs not associated with ribosomes, fraction 2 contained mRNAs associated with one or two ribosomes, fraction 3 contained mRNAs associated with 3-6 ribosomes (referred to hereafter as ‘light polysomes’), and fraction 4 contained mRNAs associated with >6 ribosomes (referred to as ‘heavy polysomes’; Fig 2A). In attached conditions, *SOD2* mRNA was primarily found in fractions 2 and 3 (Fig 2B&C), suggesting that *SOD2* is translated at a constitutive level in this condition, which is evident by our ability to readily detect SOD2 protein by western blotting. In anchorage independent conditions the relative proportion of *SOD2* mRNA shifted to fractions 3 and 4. In particular, anchorage independent cells showed a significant shift towards an enrichment of *SOD2* mRNA in the heavy polyribosome fraction 4, demonstrating a larger number of ribosomal units associated with the *SOD2* mRNA and an increase in *SOD2* mRNA translation in anchorage independent conditions. As a point of comparison, the mRNA of the nutrient stress response protein ATF4 also shifted into fraction 4 in response to anchorage-independence (Supp Fig 2).

**Figure 2.**
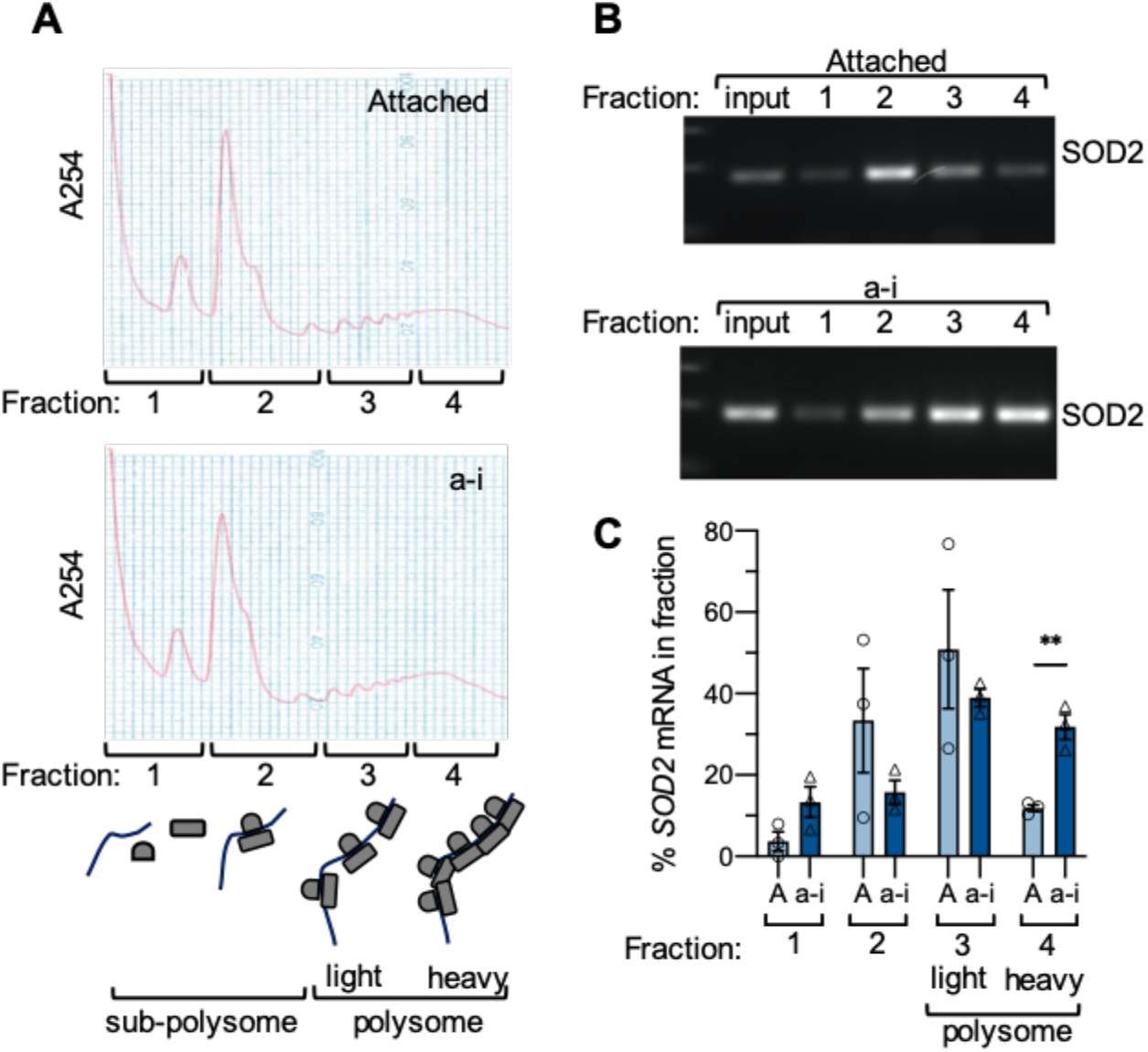
A. Polyribosome profiling was carried out after OVCA433 cells were cultured in attached (A) and anchorage independent (a-i) conditions (0.5 h) and analyzed by sucrose density gradient centrifugation. Four fractions were collected as indicated, and RNA extracted. B. Polyribosome profiling demonstrates an increase in the percentage of *SOD2* mRNA in the heavy polysomal fraction 4 in response to anchorage independence. Representative image of *SOD2* RT-PCR from RNA isolated from each polysomal fraction. C. Quantification of relative *SOD2* mRNA levels in each fraction demonstrates increased proportion of *SOD2* in fraction 4 following culture in anchorage independent conditions (*n*=3; t-test, \*\**P*<0.01).

### HuR accumulates in the cytosol and binds *SOD2* mRNA in response to anchorage-independence

Regulation of gene expression at the translational level is mediated by the interplay between mRNAs and RNA binding proteins. HuR (encoded by the gene *ELAVL1*) is a major RNA binding protein that has been implicated with alternative splicing, mRNA stability, and translation during stress conditions (Akaike *et al*, 2014; Lafarga *et al*., 2009; Lal *et al*., 2014). HuR recognizes and binds to AU-, U-rich elements in target mRNA transcripts. Screening of publicly available transcriptome-wide data sets analyzing HuR RNA binding by RNA immunoprecipitation sequencing (RIP-seq; ENCODE: ENCSR000CWW, ENCSR000CWZ) and photoactivatable ribonucleoside-enhanced crosslinking and immunoprecipitation (PAR-CLIP; GSE29943) revealed that the *SOD2* mRNA contains multiple binding sites for HuR (Fig 3A) (Davis *et al*, 2018; EncodeProjectConsortium, 2012; Lebedeva *et al*., 2011). Several clusters of HuR binding were identified in the *SOD2* 3’ UTR within 3.5 kb downstream of the STOP codon (Fig 3A). While the 5’ UTR of *SOD2* is less than 75 bp in length, the complete *SOD2* 3’ UTR spans 13,424 bp (Fig 3A, Variant 1: NM_000636). *SOD2* transcripts with variable 3’ UTR lengths have previously been reported (Suppl Fig 3A) (Chaudhuri *et al*, 2012; Church, 1990). Using RT-PCR we confirmed that OVCA433 and OVCAR10 cells express the longer 3.4 kb 3’ UTR containing the majority of HuR sites identified (Suppl Fig 3B).

**Figure 3.**
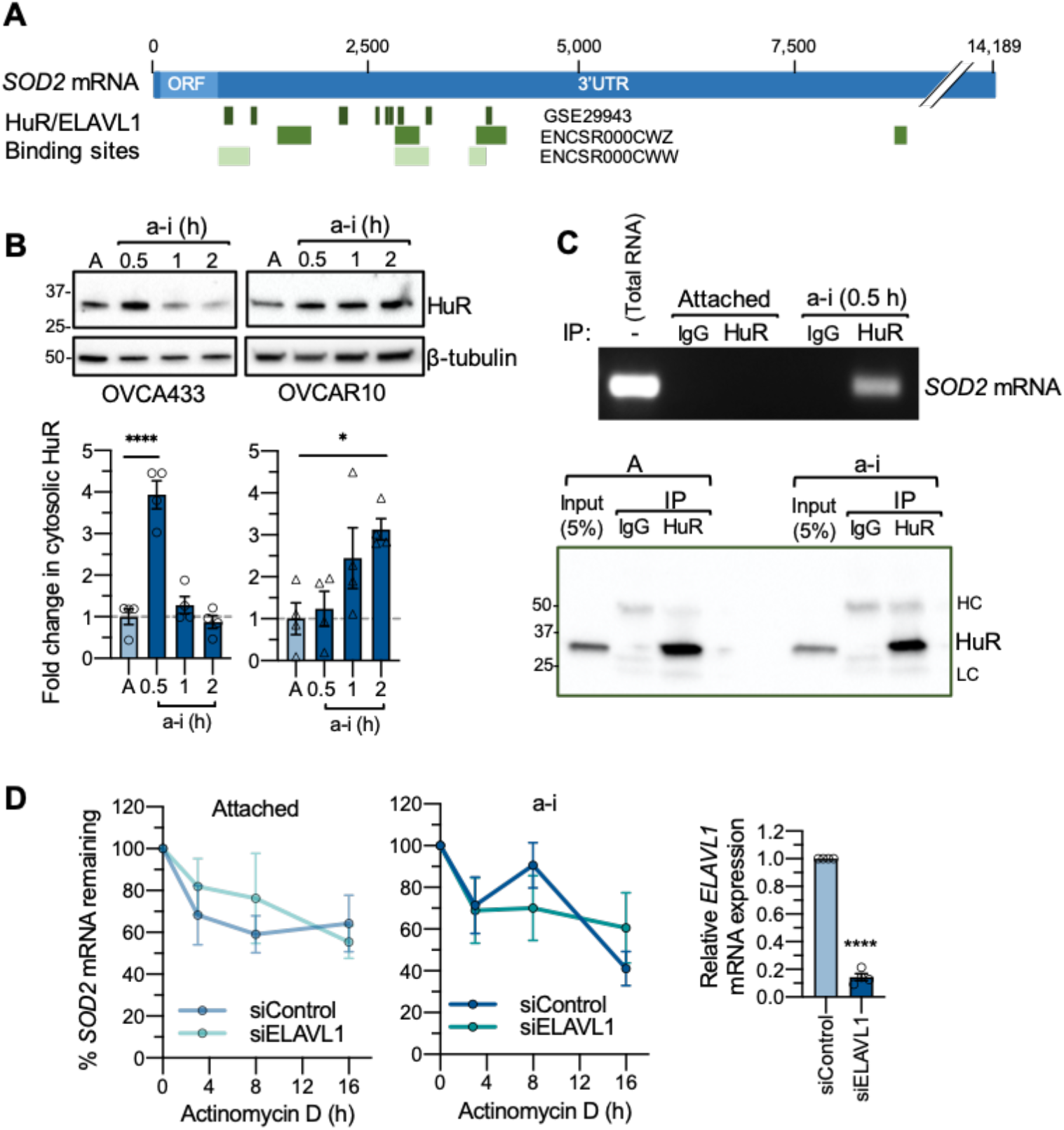
A. HuR/ELAVL1 binding profiles on the *SOD2* mRNA was assessed using ENCODE RIP-seq data sets ENCSR000CWW and ENCSR000CWZ, and PAR-CLIP data set GSE29943. B. HuR accumulates in the cytosol in response to anchorage-independence (*n*=4, one-way ANOVA, OVCA433 *P*<0.0001, OVCAR10 *P*=0.0248, Dunnett’s multiple comparison test \*\**P*<0.01; \*\*\**P*<0.001). C. Anchorage-independence induces HuR binding to *SOD2* mRNA, as assessed by Ribonucleoprotein Immunoprecipitation and *SOD2* RT-PCR following OVCA433 culture in attached or anchorage independent conditions (a-i, 0.5h). D. HuR knock-down does not affect *SOD2* mRNA stability in attached or anchorage-independent conditions, as determined by Actinomycin D treatment (*n*=4; two-way ANOVA: ns). HuR knock-down was assessed by semi quantitative real time RT-PCR (t-test, \*\*\*\**P*<0.0001).

To examine if HuR regulates SOD2 protein expression in response to anchorage independence, cytosolic translocation of HuR in response to culture in ULA plates was first determined. Concurrent with the increases in cytosolic SOD2 protein expression (Fig 1A), HuR cytosolic protein levels increased significantly in OVCA433 within 0.5 hours of anchorage independence and within 2 hours in OVCAR10 cells (Fig 3B). Based on this finding, we next investigated if HuR binds to *SOD2* mRNA in anchorage independent conditions using ribonucleoprotein immunoprecipitation to capture the HuR bound mRNAs using HuR antibody (Fig 3C). While *SOD2* mRNA could not be detected following HuR IP from attached cells, *SOD2* mRNA was readily identified by PCR in HuR immunoprecipitates from both OVCA433 (Fig 3C) and OVCAR10 cells (Supp Fig 3C) under anchorage independent conditions, indicating that matrix detachment causes the binding of HuR to *SOD2* mRNA.

Since HuR binds to *SOD2* mRNA shortly after matrix detachment, we investigated the functional consequences of the HuR-*SOD2* mRNA interaction using siRNA mediated knockdown of HuR/*ELAVL1* (Fig 3D & Fig 4A). An established function of HuR as a stress response RNA binding protein is its role in mRNA stabilization within the cytosol (Filippova *et al*., 2011; Jakstaite *et al*, 2015). To determine if HuR has an effect on *SOD2* mRNA stability, we treated ovarian cancer cells with the transcription inhibitor actinomycin D. Compared to attached conditions, anchorage independence did not significantly alter *SOD2* mRNA stability in OVCA433 cells (Fig 3D), while decreased SOD2 mRNA stability in anchorage independence was observed in OVCAR10 cells compared to attached conditions (Supp Fig 3D, two-way ANOVA, P=0.0104), indicating that these cells differ in mechanisms regulating *SOD2* mRNA stability. However, HuR knockdown did not significantly alter *SOD2* mRNA levels in response to actinomycin D treatment in anchorage independent or attached culture conditions (Fig 3D & Supp Fig 3D), suggesting that increased binding of HuR to *SOD2* mRNA does not influence *SOD2* mRNA stability.

**Figure 4.**
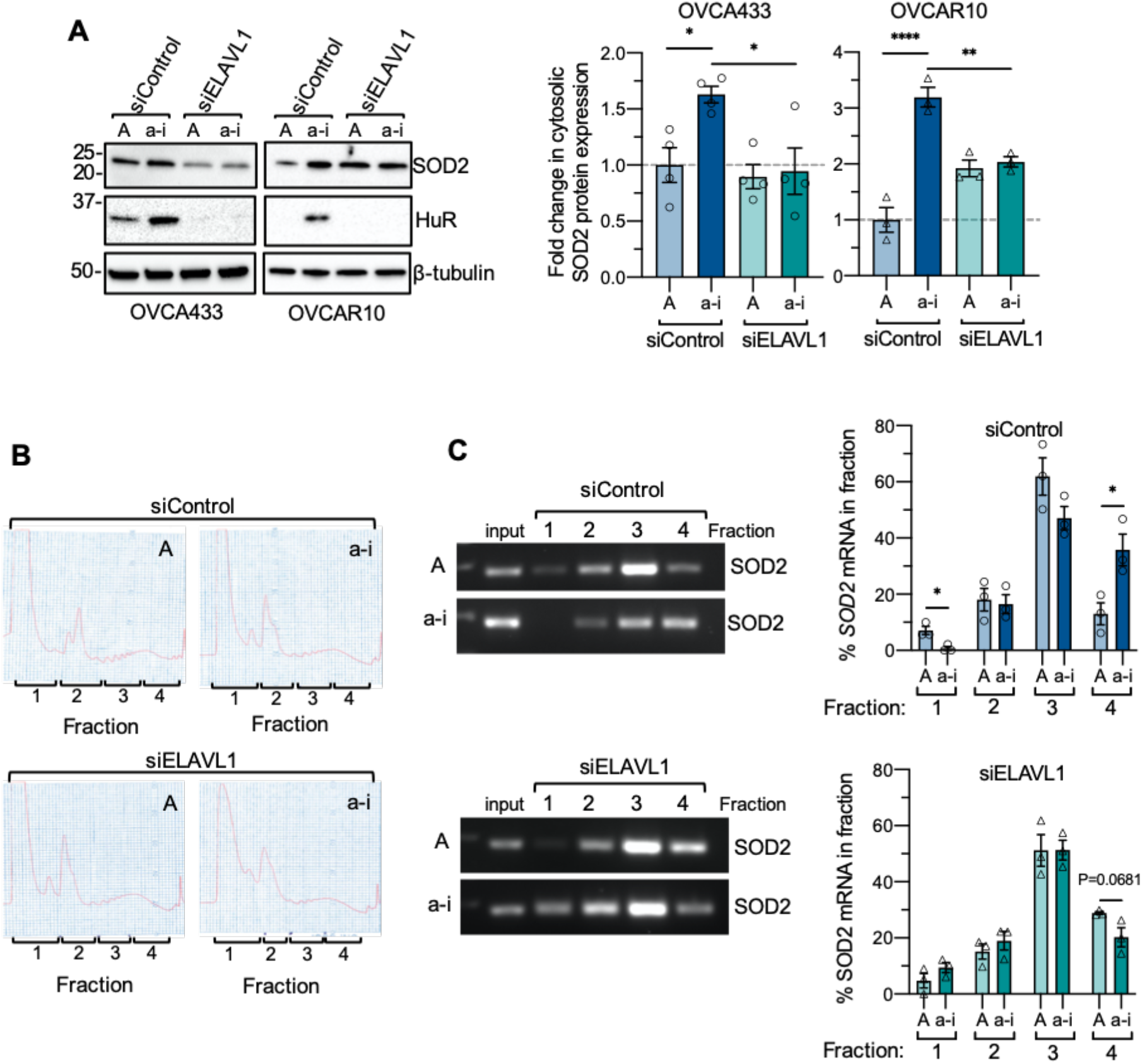
A. HuR/ELAVL1 knock-down abrogates increases in cytosolic SOD2 expression in short term anchorage-independence (a-i, OVCA433 0.5 h; OVCAR10 2 h) compared to attached cultures (A; *n*=3-4, one-way ANOVA, OVCA433 *P*=0.012, OVCAR10 *P*=0.0001; Tukey’s multiple comparison test \**P*<0.05, \*\**P*<0.01, \*\*\*\**P*<0.0001). B. Polysome profiles of OVCA433 cells cultured in attached (A) and anchorage independent (a-i, 0.5 h) conditions following siRNA-mediated HuR/ELAVL1 knockdown. C. HuR knock-down abrogates a shift of *SOD2* mRNA into fraction 4 in response to anchorage independence (a-i). Representative image of *SOD2* RT-PCR from polyribosome fractions and quantification of relative *SOD2* mRNA levels in each fraction shown (*n*=3; t-test, \**P*<0.05).

### HuR enhances *SOD2* mRNA translation under anchorage independence

We next tested if HuR is necessary for enhanced *SOD2* mRNA translation in anchorage independence. Following siRNA-mediated HuR (ELAVL1) knockdown the increase in SOD2 cytosolic protein levels induced by matrix detachment were significantly decreased (Fig 4A). To further demonstrate that increased SOD2 protein synthesis in anchorage independent ovarian cancer cells is HuR dependent, we conducted polyribosome profiling following siRNA mediated HuR knock-down (Fig 4B). In response to culture in anchorage independent conditions, *SOD2* mRNA shifted towards the heavy polyribosome fraction (fraction 4) in OVCA433 cells transfected with a scramble control siRNA (Fig 4C), as demonstrated above in un-transfected cells (Fig 2). HuR knockdown abrogated this shift of *SOD2* mRNA to the heavy polyribosomal fraction, and anchorage independent cultured cells lacking HuR displayed a similar distribution of *SOD2* mRNA in polysomal fractions compared to attached cells (Fig 4C). There was no difference in *SOD2* mRNA abundance in the subpolysome fractions (fractions 1 & 2) following HuR knock-down, indicating that a loss of HuR does not lead to a complete loss of *SOD2* mRNA translation, and suggests that the primary function of HuR is to enhance *SOD2* translation in response to anchorage independence, boosting SOD2 protein levels under these conditions.

### Inhibition of p38 MAPK activation in response to anchorage independence abrogates increases in SOD2 protein expression and HuR-*SOD2* mRNA binding

HuR can be activated in response to cellular stress *via* the p38 MAPK stress response kinase pathway (Tran *et al*, 2003; Wang *et al*., 2000b). An increase in p38 MAPK phosphorylation was previously reported in ovarian cancer cell lines cultured in anchorage independence for 24-48 h (Carduner *et al*, 2014). We were able to show that short-term anchorage independence (0.5-2 h) also increased p38 MAPK phosphorylation in OVCA433 and OVCAR10 cell lines (Fig 5A). To determine if the p38 MAPK pathway is involved in the observed increases in cytosolic SOD2 protein expression during this time, cells were treated with the p38 MAPK inhibitor, SB203580. SB203580 inhibited the phosphorylation of the p38 target MAPKAPK2 and abrogated the increases in SOD2 protein expression observed in anchorage independent conditions (Fig. 5B). In addition, the formation of the HuR-*SOD2* mRNA complex was monitored in the presence of p38 MAPK inhibition. Similar to Fig 3, anchorage independent conditions increased *SOD2* mRNA binding to HuR, while treatment with SB203580 decreased this interaction (Fig. 5C). The above demonstrates a link between p38 MAPK signaling, HuR binding to the *SOD2* mRNA and SOD2 expression in response to cellular detachment. p38 MAPK has previously been shown to phosphorylate Thr 118 of HuR (Lafarga *et al*., 2009; Liao *et al*, 2011). In the absence of a commercially available phospho-Thr118 HuR specific antibody we were unable to successfully demonstrate that anchorage independence or p38 MAPK inhibition influences phosphorylation of HuR using HuR IP and a pan phospho-Thr antibody (data not shown). In addition, we analyzed the effects of p38 MAPK on HuR cellular localization in matrix detached cells, as this had previously been shown as a mechanism of HuR regulation in response to stress (Slone *et al*, 2016; Tran *et al*., 2003; Wang *et al*., 2000b). In OVCAR10 cells, p38 MAPK inhibition decreased cytosolic HuR accumulation in response to anchorage-independence, while this could not be consistently observed in OVCA433 cells (Fig. 5D). Although it is beyond the scope of the present work, the precise mechanism by which p38 MAPK activates HuR require further attention.

**Figure 5.**
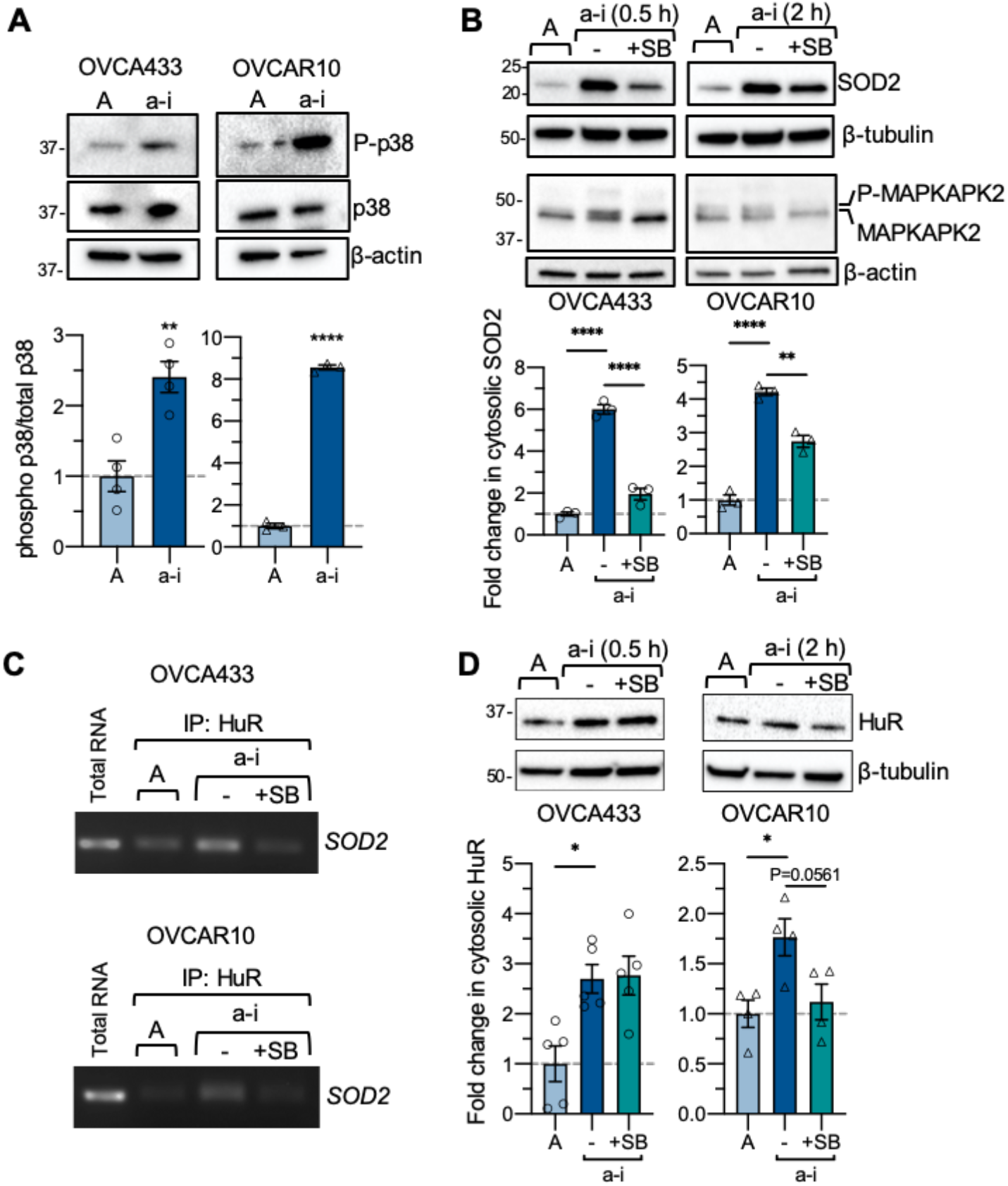
A. p38 MAPK (Thr180/Tyr182) phosphorylation is induced in response to culture in anchorage-independent culture conditions (a-i OVCA433 0.5 h, OVCAR10 2 h; n=4, T-test, \*\**P*<0.01, \*\*\*\**P*<0.0001). B. p38 MAPK inhibition abrogates a-i induced increases in SOD2 expression (n=3, one-way ANOVA *P*<0.0001, Tukey’s multiple comparison test \*\**P*<0.01, \*\*\*\**P*<0.0001). C. p38 MAPK inhibition abrogates HuR binding to *SOD2* mRNA in anchorage-independence, as assessed by RNA immunoprecipitation. D. Effects of p38 MAPK inhibition on cytosolic HuR levels (n=4-5, one-way ANOVA, OVCA433 *P*=0.0053, OVCAR10 *P*=0.0221, Tukey’s multiple comparison test \*\**P*<0.01, \*\*\*\**P*<0.0001).

The above data demonstrate a novel role for HuR in the rapid upregulation of *SOD2* translation in response to detachment of ovarian cancer cells. We previously reported that SOD2 is necessary for survival of cells in anchorage independence (Kim *et al*., 2020). Similarly, HuR knock-down resulted in an increase in the dead cell fraction of OVCA433 cells cultured for 2 h in anchorage independent conditions (Fig 6). These data demonstrate that HuR contributes to cellular survival in early stages of detachment.

**Figure 6.**
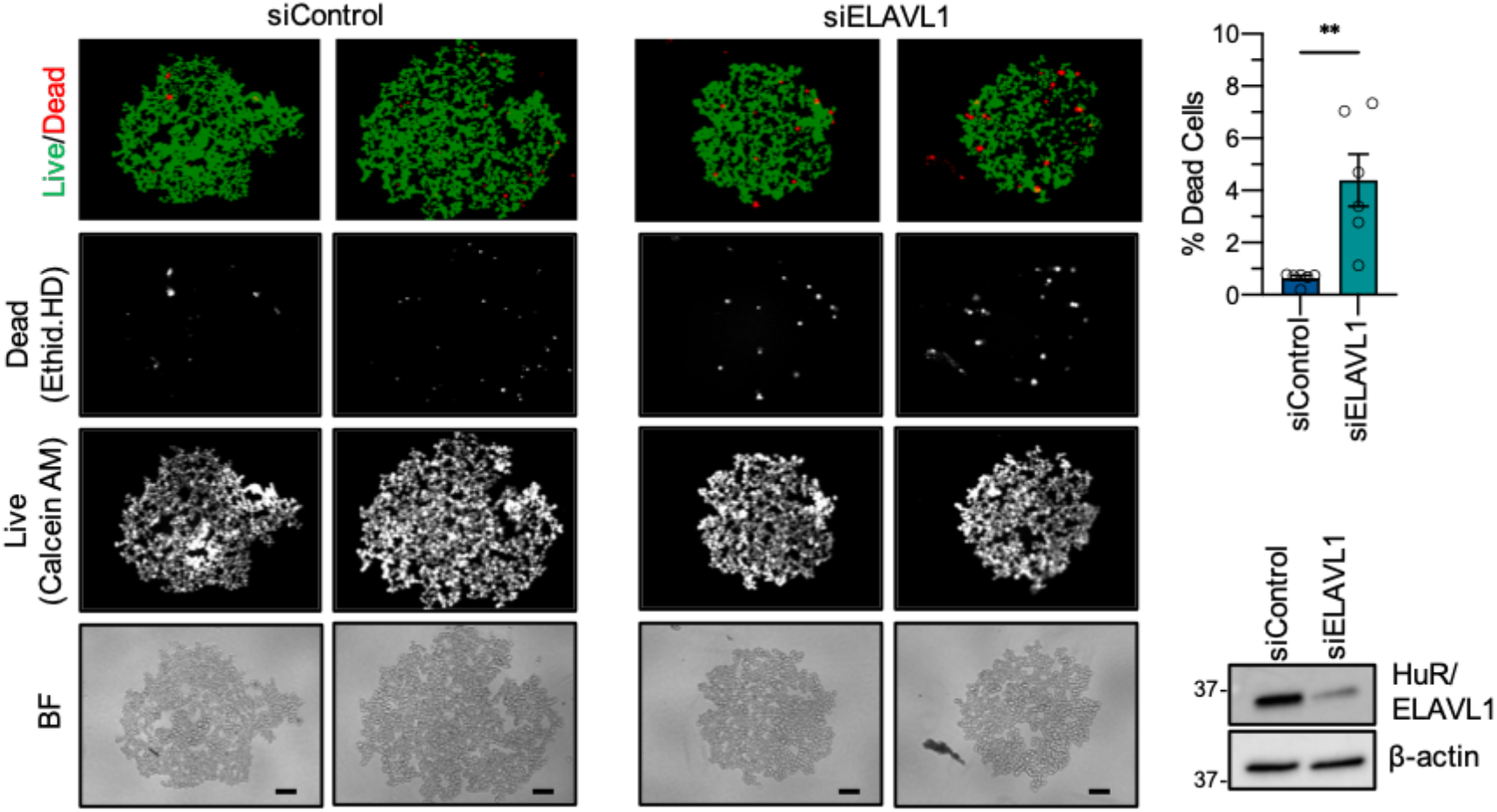
HuR/ELAVL1 knock-down increases the dead cell fraction of OVCA433 cells when cultured in anchorage-independence for 2 h. Cells were stained with Ethidium homodimer (dead cells) and Calcein AM (live cells) and fractions of live and dead cells quantified (scale bar = 100μm; n=6; T-test \*\**P*=0.004).

## Discussion

Recent studies have highlighted that tumor cells need an adequate antioxidant system to deal with intrinsic and extrinsic increases in ROS associated with metastatic progression (Davison *et al*., 2013; Kim *et al*., 2020; Piskounova *et al*., 2015). Tumor cells must therefore readily adapt to increase their antioxidant capacity at the transcriptional and post-transcriptional levels. In line with these findings, we previously showed that SIRT3-mediated deacetylation of SOD2 drives transcoelomic metastasis by increasing mitochondrial antioxidant capacity in anchorage-independent ovarian cancer cells (Kim *et al*., 2020). The present work demonstrates an additional SOD2 regulatory mechanism during early-stage anchorage independence. We found that ovarian cancer cells rapidly adapt to detachment by increasing *SOD2* mRNA translation in a p38 MAPK-HuR-dependent manner.

Aberrant HuR expression has been reported in several malignancies including ovarian cancer, and an increase in cytoplasmic subcellular localization, where HuR acts to enhance translation, is linked with worse prognosis (Denkert *et al*., 2004a; Denkert *et al*., 2004b; Miyata *et al*., 2013). HuR’s pro-tumorigenic function in the cytoplasm involves selective stabilization and/or increased translation of target mRNAs. Previously identified HuR targets not only include mRNAs encoding pro-survival and anti-apoptotic proteins, such as Bcl-2 (Filippova *et al*., 2011; Ishimaru *et al*, 2009; Wang *et al*, 2000a), but also angiogenic factors and proteins that support invasion and metastasis, such as VEGF (Levy *et al*, 1998; Tran *et al*., 2003). Similar to our findings (Fig 6), HuR knock-down decreased glioma cell survival in anchorage independence (Filippova *et al*., 2011). It was found that HuR knock-down increased apoptosis and decreased *Bcl-2* mRNA stability and protein expression, and the investigators demonstrated that HuR has the ability to bind the 3’ UTR of *Bcl-2* family members (Filippova *et al*., 2011). Moreover, HuR regulation can interplay with miRNAs to further fine tune expression in cancer, as has been demonstrated in ovarian cancer with miR-200c (Prislei *et al*, 2013). This continuously growing repertoire of cancer-related mRNAs regulated by HuR further emphasizes the critical role of this RNA binding protein in cancer cells. Moreover, HuR appears to confer survival advantages to cancer cells by rapidly adjusting gene expression for stress adaptation and resistance to cell death. Our data here identify SOD2, an important antioxidant enzyme for maintaining mitochondrial redox homeostasis, as a novel HuR target during early-stage metastasis.

HuR is a predominantly nuclear protein which translocates to the cytoplasm upon extrinsic or intrinsic stimuli and stress signals. Posttranslational modifications of HuR by different signaling pathways have been shown to affect its RNA binding affinity, nucleo-cytoplasmic shuttling, and the stability of HuR protein depending on the location of residues (Abdelmohsen & Gorospe, 2010). In particular, phosphorylation events are directly involved in spatiotemporal regulation of HuR. Among different kinases activated during stress, p38 MAPK-dependent phosphorylation on Thr118 induces cytoplasmic accumulation of HuR and increased p21 mRNA binding after exposure to ionizing radiation (Lafarga *et al*., 2009) and enhanced mRNA binding upon IL-1β treatment (Liao *et al*., 2011). Consistent with these previous findings, we found that stress associated with matrix detachment activated p38 MAPK (Fig 5). Importantly, activation of the p38 MAPK pathway increased SOD2 cytosolic protein expression under anchorage independence and we found that the association of HuR with *SOD2* mRNA was also p38 MAPK-dependent (Fig 2 & 5). It remains to be determined whether HuR is phosphorylated on Thr118 in anchorage independent cells, or if p38 MAPK indirectly activates HuR to bind *SOD2* mRNA. Moreover, cytosolic HuR accumulation was not affected by the p38 MAPK inhibitor in one of the ovarian cancer cells (Fig 5), raising a possibility that additional stress signaling pathways could contribute to the HuR nucleo-cytoplasmic shuttling in OVCA433 cells.

The p38 MAPK pathway primarily governs the cellular response to stress and triggers apoptosis upon matrix detachment or oncogene-induced ROS accumulation (Dolado *et al*, 2007; Owens *et al*, 2009). During cancer progression, however, elevated levels of intracellular oxidants accelerate proliferation and transition to malignant phenotypes via upregulation of pro-survival pathways (Liou *et al*, 2016; Weinberg *et al*, 2010), and cancer cells must activate antioxidant defenses to maintain these elevated ROS at sub-lethal levels. Cancer cells can acquire resistance by uncoupling the p38 MAPK pro-apoptotic pathway from it’s ROS-sensing ability (Dolado *et al*., 2007). Therefore, cancer cells manipulate the cellular p38 MAPK surveillance program to favor their survival and adapt to a variety of stresses that they encounter during different stages of cancer. Indeed, downstream targets of p38 MAPK, including HIF-1α and uPA (urokinase-type plasminogen activator) (Emerling *et al*, 2005; Huang *et al*, 2000) are well-known drivers of angiogenesis and cancer invasion. Here we show that increased cytosolic SOD2 protein expression was also dependent on p38 MAPK activity, demonstrating that cancer cells overcome matrix detachment-induced stress via this pathway. Further investigation of the downstream mediators linking the p38 MAPK pathway to HuR under conditions of anchorage independence require further attention and should unveil novel stress response proteins important for anoikis resistance.

While the transcriptional regulation of antioxidant enzymes has been studied widely in the context of antioxidant response elements and stress response transcription factors such as Nrf2/NFE2L2, fewer studies have focused on translational regulation of these enzymes. In earlier work, the presence of an unknown stress-responsive *SOD2* mRNA binding protein was reported across several cells and tissues of different species (Chung *et al*, 1998; Fazzone *et al*, 1993). Rat lung extracts contained a redox-sensitive *SOD2* mRNA binding protein (Fazzone *et al*., 1993), and further analysis identified that this is associated with a cis-regulatory region located 111 bp downstream of the stop codon in rat *SOD2* mRNA (Chung *et al*., 1998). The 3’ UTR of human *SOD2* mRNA shares ∼75% homology with the rat 3’ UTR. Based on sequence comparison, we found that the sequence of the protein binding region partially overlaps with the first HuR binding sites from the PAR-CLIP analysis (Fig 3A) (Chung *et al*., 1998; Lebedeva *et al*., 2011). Among the different *SOD2* mRNA splice variants, different 3’ UTRs have been reported (Supp Fig 3A). Variant 2 (NM_001024465) has a short 3’ UTR composed of a spliced region that excludes the majority of the HuR sites identified. Variant 1 (NM_000636) has been annotated to contain a 13.4 kb 3’ UTR. However, past studies have shown that the two most common *SOD2* transcripts contain either a short 240 bp or a 3,439 bp segment of this 3’ UTR, which arise from use of a proximal and distal polyadenylation site, respectively (Supp Fig 3A) (Chaudhuri *et al*., 2012; Church, 1990). Interestingly, Chaudhuri *et al*. reported that the expression of these two *SOD2* transcripts is altered between quiescent and proliferating cells, with the shorter transcript being associated with quiescence and increased protein expression (Chaudhuri *et al*., 2012). Moreover, radiation increased levels of the shorter *SOD2* transcript levels of the 1.5 kb MnSOD transcript, with expression of the longer form remaining unaltered (Chaudhuri *et al*., 2012). The mechanisms for this radiation induced increase in the short 3’ UTR transcript remain unclear. However, we predict that is likely not HuR-dependent, as only the longer 3.4 kb 3’ UTR contains the majority of identified HuR binding sites, and we verified that ovarian cancer cells used in the present work express the transcript containing this longer 3’ UTR (Supp Fig 3B). It remains to be determined if and how these alternate 3’ UTR *SOD2* transcripts are regulated during different sources of stress, and how their transcription co-operates with translational regulation through the activation of cell-specific RNA binding proteins, as well as the interplay with non-coding RNAs, such as miRNAs. A screen for miRNA binding reveals that the *SOD2* mRNA contains potential binding sites for miRNAs throughout the length of the 3’ UTR. While most are located toward the far upstream region, several overlap with identified HuR binding sites. The role of miRNAs in regulating SOD2 expression have also been interrogated and several miRNAs identified that either positively or negatively regulate SOD2 levels in cancer (Kim *et al*., 2017). It remains to be determined if changes in miRNA binding further influence the regulation of *SOD2* mRNA translation in anchorage-independence, and if this interplays with the regulation by HuR.

In conclusion, we show for the first time that *SOD2* mRNA is an HuR target in anchorage-independent ovarian cancer cells. Although SOD2 regulatory mechanisms have been studied extensively at the transcriptional level, very few studies have investigated how translational regulation contributes to adaptable changes in SOD2 expression. The present findings uncover a novel post-transcriptional stress response mechanism by which tumor cells are able to rapidly increase their mitochondrial antioxidant capacity to adapt to stress associated with anchorage-independence, promoting survival during metastatic progression.

## Materials and Methods

### Cell Culture and Reagents

OVCA433 and OVCAR10 cells were provided by Dr. Susan K. Murphy (Duke University) and Dr. Katherine Aird (University of Pittsburgh), respectively. OVCA433 and OVCAR10 were grown in RPMI1640 supplemented with 10% FBS at 37 °C with 5% CO_2_. STR profiling is carried out routinely to validate cell identity, which revealed at the commencement of this work that OVCAR10 cells share the same STR profile as NIH-OVCAR3 cells. It is unclear if the OVCAR10 cell line was initially derived from the same patient as OVCAR3, or if OVCAR10 cells represent a sub-line derived from OVCAR3 cells. The protein synthesis inhibitor cycloheximide (Sigma) was added at a concentration of 20 µg/mL in fully supplemented growth media. For mRNA stability assays, actinomycin D (Sigma) was added at 10 µg/mL. The p38 MAPK inhibitor SB203580 was used at a final concentration of 20 µM.

### Cell culture in adherent and ultra-low attachment (ULA) conditions

For attached conditions, cells were plated in 150-mm dishes and grown to ∼80% confluency. For anchorage independent cell culture, cells were trypsinized and seeded at a density (300,000 cells/2 mL media/well) in 6-well ULA (ultra-low attachment) plates (Corning: 3471) and collected at different time points for downstream analyses.

### siRNA-mediated HuR/ELAVL1 knock-down

Cells were transfected with scramble non-targeting SMARTpool control (Dharmacon: D-001810-10-05) or HuR (ELAVL1)-specific SMARTpool siRNA oligonucleotides (Dharmacon: L-003773-00-0005) using Lipofectamine RNAiMAX (Invitrogen), and knock-down confirmed by western blotting.

### Subcellular Fractionation

Cells in adherent and ULA plates were collected and the cell pellets were washed with ice-cold PBS. The cell pellets were processed as described in Sugiura *et al*. (Sugiura *et al*, 2017). Briefly, cells were centrifuged and resuspended in 200-500 µl of ice-cold homogenization buffer (10 mM HEPES pH 7.4, 220 mM mannitol, 70 mM sucrose, Roche protease and phosphatase inhibitor cocktails). The lysates were homogenized by several passages through 27-G needles. Lysates were centrifuged at 800 g for 10 min, followed by centrifugation of the supernatants at 2,500 g for 15 min at 4 °C. The mitochondrial pellets were resuspended in homogenization buffer and the supernatants were centrifuged at 100,000g for 1 h at 4 °C using a Beckman Coulter Optima MAX Ultracentrifuge. Post-centrifugation supernatants containing cytosolic fractions were transferred to new tubes and used for immunoblotting.

### Immunoblotting

Protein concentrations were measured using the Pierce BCA protein assay kit. An equal amount of protein lysates was loaded onto 4-20% SDS-PAGE gels. Following electrophoresis, proteins were transferred to PVDF membranes. For detection of proteins, the membranes were incubated with the following antibodies overnight at 4 °C: SOD2 (A-2, Santa Cruz: sc-133134, 1:500 dilution); β-tubulin (9F3, Cell Signaling Technology: 2128, 1:1,000 dilution), ATP5A (Abcam: ab14748, 1:1000 dilution), β-actin (Thermo: AM4302, 1:10,000 dilution), HuR/ELAVL1 (3A2, Santa Cruz: sc-5261, 1:500 dilution), Phospho-p38 MAPK (Thr180/Tyr182, Cell Signaling Technology: 9211, 1:1000 dilution), p38 MAPK (A-12, Santa Cruz Biotechnology: sc-7972, 1:1000 dilution), MAPKAPK-2 (Cell signaling technology: 3042, 1:1000 dilution). The blots were developed using SuperSignal West Femto Maximum Sensitivity Substrate (Thermo: 34096) after incubation with horseradish peroxidase (HRP)-conjugated secondary antibodies (Amersham Biosciences) for 1 h at RT.

### Immunoprecipitation (IP)

1-1.5 mg of cell lysates were pre-cleared by incubating with 2 µg normal rabbit IgG (Cell Signaling Technology: 2729S) or normal mouse IgG (Millipore: 12-371) on a rotator for 1 h at 4 °C followed by an additional 1 h incubation with protein A-(Thermo: 20333) or protein G-agarose beads (50 µL; Thermo: 20399) at 4 °C. Following centrifugation at 3000g for 10 min supernatants were transferred to clean tubes and incubated with either IgG or primary antibodies overnight at 4 °C. 50 µL of agarose beads were added to the lysates for 1-2 h at 4 °C and the antibody-bead complexes were washed three times in IP lysis buffer and further processed for downstream assays.

### ^35^S Protein Radiolabeling

Cells in adherent and ULA plates were treated with EasyTag Express^35^S Protein Labeling Mix (Perkin Elmer: NEG772), using 40 µl ^35^S (440 uCi) per 20 mL media in 150-mm dish, 4 µl ^35^S (44 uCi) /2 mL media/ well in ULA plates, according to a protocol adapted from Gallagher *et al*. (Gallagher *et al*, 2008). Following 2 h incubation in the presence of ^35^S-L-methionine and ^35^S-L-cysteine, cells were collected, washed with ice-cold PBS, and harvested using RIPA buffer supplemented with protease and phosphatase inhibitors. The cell lysates were rotated for 30 min at 4 °C, centrifuged at 12,000 rpm for 30 min at 4 °C and supernatants transferred to new tubes. After pre-clearing, the lysates were incubated overnight with 2 µg of normal rabbit IgG or SOD2 antibody (Abcam: Ab13533). Following SOD2 IP, the lysates were resolved in SDS-PAGE gels. The SOD2 band in each lane was cut with a razor blade and weighed. The bands were dissolved in 1 mL of 1X TGS running buffer overnight on a rocker at 4 °C. Next day, dissolved gel pieces were further heated for 20 min at 60 °C. The dissolved radioactive sample solutions were transferred to glass vials containing 10 mL of Opti-Fluor (Perkin Elmer) in duplicate (500 µl per vial). Liquid scintillation counting was performed using a Beckman Coulter Scintillation Counter. The readouts were normalized against the values from untreated samples.

### Ribonucleoprotein Immunoprecipitation & RT-PCR

Cells were cultured in attached and anchorage independent conditions as described above. Before harvesting cells, 0.3% formaldehyde was added for 10 min at 37 °C for crosslinking followed by addition of glycine (final concentration 0.25 M) for 5 min for quenching. RNP-IP was performed as described in (Raspaglio *et al*., 2010; Tenenbaum *et al*, 2002) with modifications. Briefly, crosslinked cells were lysed in 500-1,000 µl NT1 buffer (100 mM KCl, 5 mM MgCl_2_, 10 mM HEPES, [pH 7.0], 0.5% Nonidet P40 [NP40], 1 mM dithiothreitol [DTT], 100 units/mL SUPERase·In RNase Inhibitor [Invitrogen: AM2694], protease inhibitors [Thermo: 78429], 0.2% vanadyl ribonucleoside complexes [New England Biolabs: S1402S]). After centrifugation of lysates at 16,000 g for 15 min, the supernatants were used for IP with normal mouse IgG or HuR antibody. The antibody-bead mixtures were washed several times with NT2 buffer (50 mM Tris-HCl [pH 7.4], 150 mM NaCl, 1 mM MgCl_2_, 0.05% NP40, RNAse inhibitor, protease inhibitor). IP samples for RNA elution were incubated with proteinase K (30 µg/100 µl NT2 buffer with 0.1% SDS) for 30 min at 60 °C. RNA was extracted using TRIzol, followed by cDNA synthesis (Quantabio: 95047) and *SOD2* RT-PCR using the PrimeSTAR polymerase (Takara: R010A) with the following cycles: 98°C for 10 sec, 98°C for 10 sec + 60°C for 10 sec + 72°C for 20 sec X 35-38 cycles, followed by a final extension step at 72°C for 2 min. PCR products were analyzed by 2% agarose gel electrophoresis.

### Polysome Profiling by Sucrose Density Gradient Centrifugation

Cells in adherent and ULA plates were incubated with cycloheximide (100 µg/mL) for 10 min at 37 °C before harvesting and were washed twice with ice cold 1X PBS containing cycloheximide. The cells were homogenized in 500 µl lysis buffer (50 mM HEPES, 75 mM KCl, 5 mM MgCl_2_, 250 mM sucrose, 100 ug/mL cycloheximide, 2mM DTT, 20 U/µl SUPERase·In RNase Inhibitor [Invitrogen: AM2694], 10% Triton X-100, 13% NaDOC) and polysome profiling carried out as previously described (Dang Do *et al*, 2009). Lysates were placed on ice for 10 min and centrifuged at 3000 g for 15 min at 4 °C. 500 µl supernatants were loaded on linear sucrose gradients ranging from 20% to 47% (10 mM HEPES, KCl 75 mM, 5 mM MgCl_2_, 0.5 mM EDTA) and were separated by ultracentrifugation in a SW41 rotor at 34,000 rpm for 4 h 15 min at 4 °C (Beckman Coulter). Subsequently, four sucrose fractions were collected using a UV/VIS absorbance detector. TRIzol reagent (Invitrogen) was added to each fraction for RNA isolation. Briefly, post-centrifugation at 3,200g for 20 min after addition of 1/5 volume of chloroform, the aqueous layer was transferred, and 1/2 volume of isopropanol was added for overnight precipitation at −20 °C. RNA was pelleted by centrifugation at 4,640 rpm for 55 min at 4 °C. RNA pellets were washed with 70% ethanol twice and dissolved in RNAse-free water. After cDNA synthesis and qPCR reactions, final PCR products were analyzed on 2% agarose gels.

### Semi-quantitative real-time PCR

Total RNA was isolated by RNA isolation kit (Zymo Research: R2052) and used for cDNA synthesis (Quantabio: 95047) according to the manufacturer’s instruction. cDNA was mixed with iTaq™ Universal SYBR® Green Supermix (BioRad) and the primers listed in Table 1. Semi-quantitative real time RT-PCR was carried out using a BioRad qRT-PCR machine (BioRad), data normalized to the geometric mean of four housekeeping genes (Table 1), and expressed as fold-change in expression using the 2^-ΔΔCT^ formula.

**Table 1:**
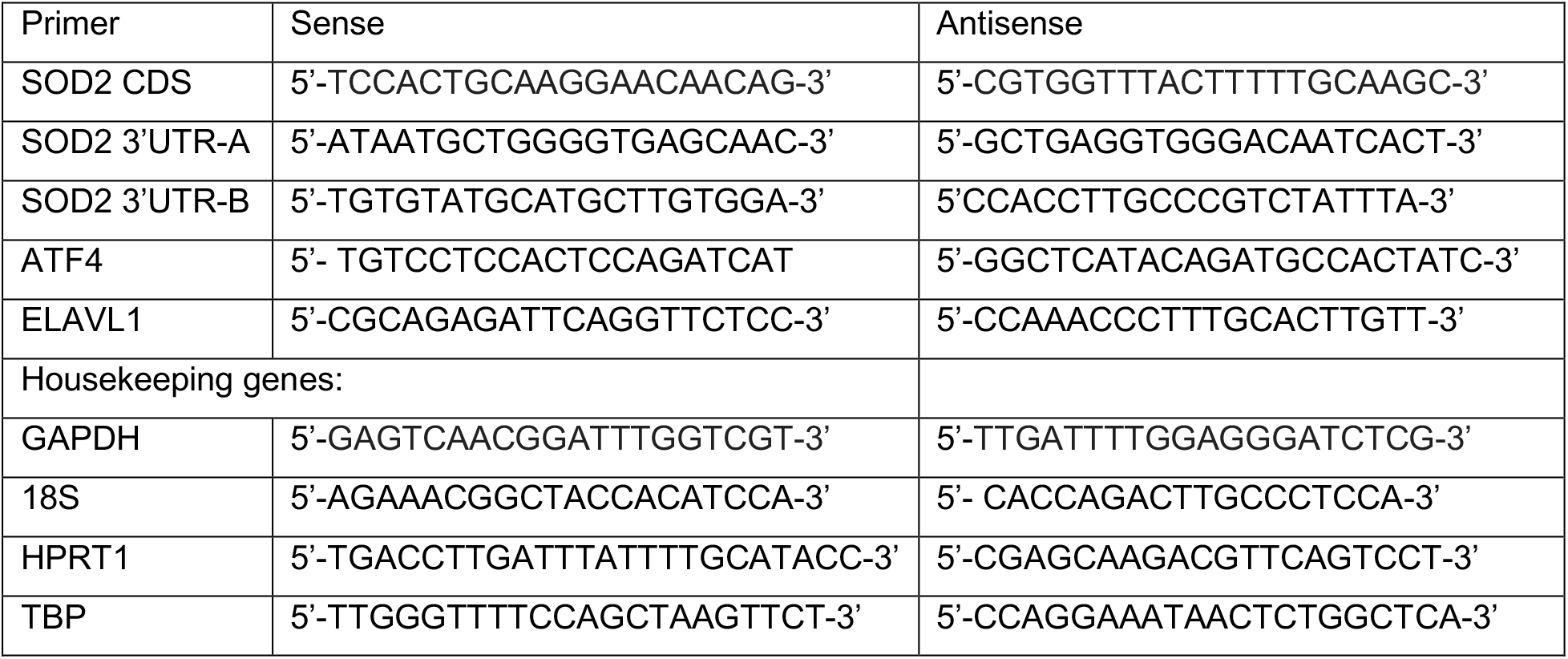
Primers used for RT-PCR and semi-quantitative real time PCR.

### Live/dead staining

Live and dead cell fractions of cells cultured for 2 h in anchorage independence was assessed by staining with 4 μM Calcein AM and 4 μM ethidium homodimer (in PBS; Sigma) to visualize live and dead cells, respectively. Cells were exposed to both dyes for 30 min at 37 °C, followed by imaging on a Keyence BZ-X700 fluorescence microscope. The percentage of live and dead cells were quantified using Image J.

### Statistical Analysis

All data are representatives of at least three independent experiments. Data are presented as mean ± SEM with individual replicate values superimposed. Statistical analysis was performed using GraphPad Prism Software v9, with statistical tests chosen based on experimental design, as described in figure legends.

## Acknowledgements

The authors would like to thank Ms. Sara Shimko and Lydia Kutzler for technical assistance. This work was supported by the U.S. National Institutes of Health grants R01CA242021 (N.H.) and R01CA230628 (N.H. & K.M.).

## Author contributions

Y.S.K. designed the conceptual framework and experiments of the study, carried out the majority of the experiments and data analysis, prepared the figures and wrote the manuscript. J.E.W., P.T., Z.J. and A.E. assisted with experimental execution, and manuscript editing. K.M. and S.R.K. contributed to experimental design, data interpretation and manuscript editing. N.H. supervised and conceived the study, contributed to experimental design, assisted in data analysis, and assisted in writing and editing of the manuscript.

## References

Abdelmohsen K, Gorospe M (2010) Posttranscriptional regulation of cancer traits by HuR. Wiley Interdiscip Rev RNA 1: 214–229

Ahmed N, Stenvers KL (2013) Getting to know ovarian cancer ascites: opportunities for targeted therapy-based translational research. Front Oncol 3: 256

Akaike Y, Masuda K, Kuwano Y, Nishida K, Kajita K, Kurokawa K, Satake Y, Shoda K, Imoto I, Rokutan K (2014) HuR regulates alternative splicing of the TRA2beta gene in human colon cancer cells under oxidative stress. Mol Cell Biol 34: 2857–2873

Audic Y, Hartley RS (2004) Post-transcriptional regulation in cancer. Biol Cell 96: 479–498

Carduner L, Picot CR, Leroy-Dudal J, Blay L, Kellouche S, Carreiras F (2014) Cell cycle arrest or survival signaling through alphav integrins, activation of PKC and ERK1/2 lead to anoikis resistance of ovarian cancer spheroids. Exp Cell Res 320: 329–342

Chaudhuri L, Nicholson AM, Kalen AL, Goswami PC (2012) Preferential selection of MnSOD transcripts in proliferating normal and cancer cells. Oncogene 31: 1207–1216

Chung DJ, Wright AE, Clerch LB (1998) The 3’ untranslated region of manganese superoxide dismutase RNA contains a translational enhancer element. Biochemistry 37: 16298–16306

Church SL (1990) Manganese superoxide dismutase: nucleotide and deduced amino acid sequence of a cDNA encoding a new human transcript. Biochim Biophys Acta 1087: 250–252

Dang Do AN, Kimball SR, Cavener DR, Jefferson LS (2009) eIF2alpha kinases GCN2 and PERK modulate transcription and translation of distinct sets of mRNAs in mouse liver. Physiol Genomics 38: 328–341

Davis CA, Hitz BC, Sloan CA, Chan ET, Davidson JM, Gabdank I, Hilton JA, Jain K, Baymuradov UK, Narayanan AK et al (2018) The Encyclopedia of DNA elements (ENCODE): data portal update. Nucleic Acids Res 46: D794–D801

Davison CA, Durbin SM, Thau MR, Zellmer VR, Chapman SE, Diener J, Wathen C, Leevy WM, Schafer ZT (2013) Antioxidant enzymes mediate survival of breast cancer cells deprived of extracellular matrix. Cancer Res 73: 3704–3715

Denkert C, Weichert W, Pest S, Koch I, Licht D, Köbel M, Reles A, Sehouli J, Dietel M, Hauptmann S (2004a) Overexpression of the embryonic-lethal abnormal vision-like protein HuR in ovarian carcinoma is a prognostic factor and is associated with increased cyclooxygenase 2 expression. Cancer Res 64: 189–195

Denkert C, Weichert W, Winzer KJ, Müller BM, Noske A, Niesporek S, Kristiansen G, Guski H, Dietel M, Hauptmann S (2004b) Expression of the ELAV-like protein HuR is associated with higher tumor grade and increased cyclooxygenase-2 expression in human breast carcinoma. Clin Cancer Res 10: 5580–5586

Dolado I, Swat A, Ajenjo N, De Vita G, Cuadrado A, Nebreda AR (2007) p38alpha MAP kinase as a sensor of reactive oxygen species in tumorigenesis. Cancer Cell 11: 191–205

Emerling BM, Platanias LC, Black E, Nebreda AR, Davis RJ, Chandel NS (2005) Mitochondrial reactive oxygen species activation of p38 mitogen-activated protein kinase is required for hypoxia signaling. Mol Cell Biol 25: 4853–4862

EncodeProjectConsortium (2012) An integrated encyclopedia of DNA elements in the human genome. Nature 489: 57–74

Epis MR, Barker A, Giles KM, Beveridge DJ, Leedman PJ (2011) The RNA-binding protein HuR opposes the repression of ERBB-2 gene expression by microRNA miR-331-3p in prostate cancer cells. J Biol Chem 286: 41442–41454

Fazzone H, Wangner A, Clerch LB (1993) Rat lung contains a developmentally regulated manganese superoxide dismutase mRNA-binding protein. J Clin Invest 92: 1278–1281

Filippova N, Yang X, Wang Y, Gillespie GY, Langford C, King PH, Wheeler C, Nabors LB (2011) The RNA-binding protein HuR promotes glioma growth and treatment resistance. Mol Cancer Res 9: 648–659

Gallagher JW, Kubica N, Kimball SR, Jefferson LS (2008) Reduced eukaryotic initiation factor 2Bepsilon-subunit expression suppresses the transformed phenotype of cells overexpressing the protein. Cancer Res 68: 8752–8760

Hart PC, Ratti BA, Mao M, Ansenberger-Fricano K, Shajahan-Haq AN, Tyner AL, Minshall RD, Bonini MG (2016) Caveolin-1 regulates cancer cell metabolism via scavenging Nrf2 and suppressing MnSOD-driven glycolysis. Oncotarget 7: 308–322

Hemachandra LP, Shin DH, Dier U, Iuliano JN, Engelberth SA, Uusitalo LM, Murphy SK, Hempel N (2015) Mitochondrial Superoxide Dismutase Has a Protumorigenic Role in Ovarian Clear Cell Carcinoma. Cancer Res 75: 4973–4984

Huang S, New L, Pan Z, Han J, Nemerow GR (2000) Urokinase plasminogen activator/urokinase-specific surface receptor expression and matrix invasion by breast cancer cells requires constitutive p38alpha mitogen-activated protein kinase activity. J Biol Chem 275: 12266–12272

Ishimaru D, Ramalingam S, Sengupta TK, Bandyopadhyay S, Dellis S, Tholanikunnel BG, Fernandes DJ, Spicer EK (2009) Regulation of Bcl-2 expression by HuR in HL60 leukemia cells and A431 carcinoma cells. Mol Cancer Res 7: 1354–1366

Jakstaite A, Maziukiene A, Silkuniene G, Kmieliute K, Gulbinas A, Dambrauskas Z (2015) HuR mediated post-transcriptional regulation as a new potential adjuvant therapeutic target in chemotherapy for pancreatic cancer. World J Gastroenterol 21: 13004–13019

Jiang L, Shestov AA, Swain P, Yang C, Parker SJ, Wang QA, Terada LS, Adams ND, McCabe MT, Pietrak B et al (2016) Reductive carboxylation supports redox homeostasis during anchorage-independent growth. Nature 532: 255–258

Kamarajugadda S, Cai Q, Chen H, Nayak S, Zhu J, He M, Jin Y, Zhang Y, Ai L, Martin SS et al (2013) Manganese superoxide dismutase promotes anoikis resistance and tumor metastasis. Cell Death Dis 4: e504

Kenny TC, Hart P, Ragazzi M, Sersinghe M, Chipuk J, Sagar MAK, Eliceiri KW, LaFramboise T, Grandhi S, Santos J et al (2017) Selected mitochondrial DNA landscapes activate the SIRT3 axis of the UPR(mt) to promote metastasis. Oncogene 36: 4393–4404

Kim YS, Gupta Vallur P, Jones VM, Worley BL, Shimko S, Shin DH, Crawford LC, Chen CW, Aird KM, Abraham T et al (2020) Context-dependent activation of SIRT3 is necessary for anchorage-independent survival and metastasis of ovarian cancer cells. Oncogene 39: 1619– 1633

Kim YS, Gupta Vallur P, Phaeton R, Mythreye K, Hempel N (2017) Insights into the Dichotomous Regulation of SOD2 in Cancer. Antioxidants (Basel) 6

Konstantinopoulos PA, Spentzos D, Fountzilas E, Francoeur N, Sanisetty S, Grammatikos AP, Hecht JL, Cannistra SA (2011) Keap1 mutations and Nrf2 pathway activation in epithelial ovarian cancer. Cancer Res 71: 5081–5089

Lafarga V, Cuadrado A, Lopez de Silanes I, Bengoechea R, Fernandez-Capetillo O, Nebreda AR (2009) p38 Mitogen-activated protein kinase- and HuR-dependent stabilization of p21(Cip1) mRNA mediates the G(1)/S checkpoint. Mol Cell Biol 29: 4341–4351

Lal S, Burkhart RA, Beeharry N, Bhattacharjee V, Londin ER, Cozzitorto JA, Romeo C, Jimbo M, Norris ZA, Yeo CJ et al (2014) HuR posttranscriptionally regulates WEE1: implications for the DNA damage response in pancreatic cancer cells. Cancer Res 74: 1128–1140

Lebedeva S, Jens M, Theil K, Schwanhäusser B, Selbach M, Landthaler M, Rajewsky N (2011) Transcriptome-wide analysis of regulatory interactions of the RNA-binding protein HuR. Mol Cell 43: 340–352

Levy NS, Chung S, Furneaux H, Levy AP (1998) Hypoxic stabilization of vascular endothelial growth factor mRNA by the RNA-binding protein HuR. J Biol Chem 273: 6417–6423

Liao WL, Wang WC, Chang WC, Tseng JT (2011) The RNA-binding protein HuR stabilizes cytosolic phospholipase A2α mRNA under interleukin-1β treatment in non-small cell lung cancer A549 Cells. J Biol Chem 286: 35499–35508

Liou GY, Döppler H, DelGiorno KE, Zhang L, Leitges M, Crawford HC, Murphy MP, Storz P (2016) Mutant KRas-Induced Mitochondrial Oxidative Stress in Acinar Cells Upregulates EGFR Signaling to Drive Formation of Pancreatic Precancerous Lesions. Cell Rep 14: 2325–2336

Mazan-Mamczarz K, Hagner PR, Corl S, Srikantan S, Wood WH, Becker KG, Gorospe M, Keene JD, Levenson AS, Gartenhaus RB (2008) Post-transcriptional gene regulation by HuR promotes a more tumorigenic phenotype. Oncogene 27: 6151–6163

Miyata Y, Watanabe S, Sagara Y, Mitsunari K, Matsuo T, Ohba K, Sakai H (2013) High expression of HuR in cytoplasm, but not nuclei, is associated with malignant aggressiveness and prognosis in bladder cancer. PLoS One 8: e59095

Mrena J, Wiksten JP, Thiel A, Kokkola A, Pohjola L, Lundin J, Nordling S, Ristimäki A, Haglund C (2005) Cyclooxygenase-2 is an independent prognostic factor in gastric cancer and its expression is regulated by the messenger RNA stability factor HuR. Clin Cancer Res 11: 7362– 7368

Owens TW, Valentijn AJ, Upton JP, Keeble J, Zhang L, Lindsay J, Zouq NK, Gilmore AP (2009) Apoptosis commitment and activation of mitochondrial Bax during anoikis is regulated by p38MAPK. Cell Death Differ 16: 1551–1562

Piskounova E, Agathocleous M, Murphy MM, Hu Z, Huddlestun SE, Zhao Z, Leitch AM, Johnson TM, DeBerardinis RJ, Morrison SJ (2015) Oxidative stress inhibits distant metastasis by human melanoma cells. Nature 527: 186–191

Prislei S, Martinelli E, Mariani M, Raspaglio G, Sieber S, Ferrandina G, Shahabi S, Scambia G, Ferlini C (2013) MiR-200c and HuR in ovarian cancer. BMC Cancer 13: 72

Raspaglio G, De Maria I, Filippetti F, Martinelli E, Zannoni GF, Prislei S, Ferrandina G, Shahabi S, Scambia G, Ferlini C (2010) HuR regulates beta-tubulin isotype expression in ovarian cancer. Cancer Res 70: 5891–5900

Schafer ZT, Grassian AR, Song L, Jiang Z, Gerhart-Hines Z, Irie HY, Gao S, Puigserver P, Brugge JS (2009) Antioxidant and oncogene rescue of metabolic defects caused by loss of matrix attachment. Nature 461: 109–113

Slone S, Anthony SR, Wu X, Benoit JB, Aube J, Xu L, Tranter M (2016) Activation of HuR downstream of p38 MAPK promotes cardiomyocyte hypertrophy. Cell Signal 28: 1735–1741

Sugiura A, Mattie S, Prudent J, McBride HM (2017) Newly born peroxisomes are a hybrid of mitochondrial and ER-derived pre-peroxisomes. Nature 542: 251–254

Tenenbaum SA, Lager PJ, Carson CC, Keene JD (2002) Ribonomics: identifying mRNA subsets in mRNP complexes using antibodies to RNA-binding proteins and genomic arrays. Methods 26: 191–198

Torre LA, Trabert B, DeSantis CE, Miller KD, Samimi G, Runowicz CD, Gaudet MM, Jemal A, Siegel RL (2018) Ovarian cancer statistics, 2018. CA Cancer J Clin 68: 284–296

Tran H, Maurer F, Nagamine Y (2003) Stabilization of urokinase and urokinase receptor mRNAs by HuR is linked to its cytoplasmic accumulation induced by activated mitogen-activated protein kinase-activated protein kinase 2. Mol Cell Biol 23: 7177–7188

van Kouwenhove M, Kedde M, Agami R (2011) MicroRNA regulation by RNA-binding proteins and its implications for cancer. Nat Rev Cancer 11: 644–656

Wang J, Guo Y, Chu H, Guan Y, Bi J, Wang B (2013) Multiple functions of the RNA-binding protein HuR in cancer progression, treatment responses and prognosis. Int J Mol Sci 14: 10015– 10041

Wang W, Caldwell MC, Lin S, Furneaux H, Gorospe M (2000a) HuR regulates cyclin A and cyclin B1 mRNA stability during cell proliferation. EMBO J 19: 2340–2350

Wang W, Furneaux H, Cheng H, Caldwell MC, Hutter D, Liu Y, Holbrook N, Gorospe M (2000b) HuR regulates p21 mRNA stabilization by UV light. Mol Cell Biol 20: 760–769

Weinberg F, Hamanaka R, Wheaton WW, Weinberg S, Joseph J, Lopez M, Kalyanaraman B, Mutlu GM, Budinger GR, Chandel NS (2010) Mitochondrial metabolism and ROS generation are essential for Kras-mediated tumorigenicity. Proc Natl Acad Sci U S A 107: 8788–8793

Wurth L, Gebauer F (2015) RNA-binding proteins, multifaceted translational regulators in cancer. Biochim Biophys Acta 1849: 881–886

