## Supplemental Figures for "HuR -dependent SOD2 protein synthesis is an early adaptation of ovarian cancer cells in response to anchorage independence"

Yeon Soo Kim *et al*.

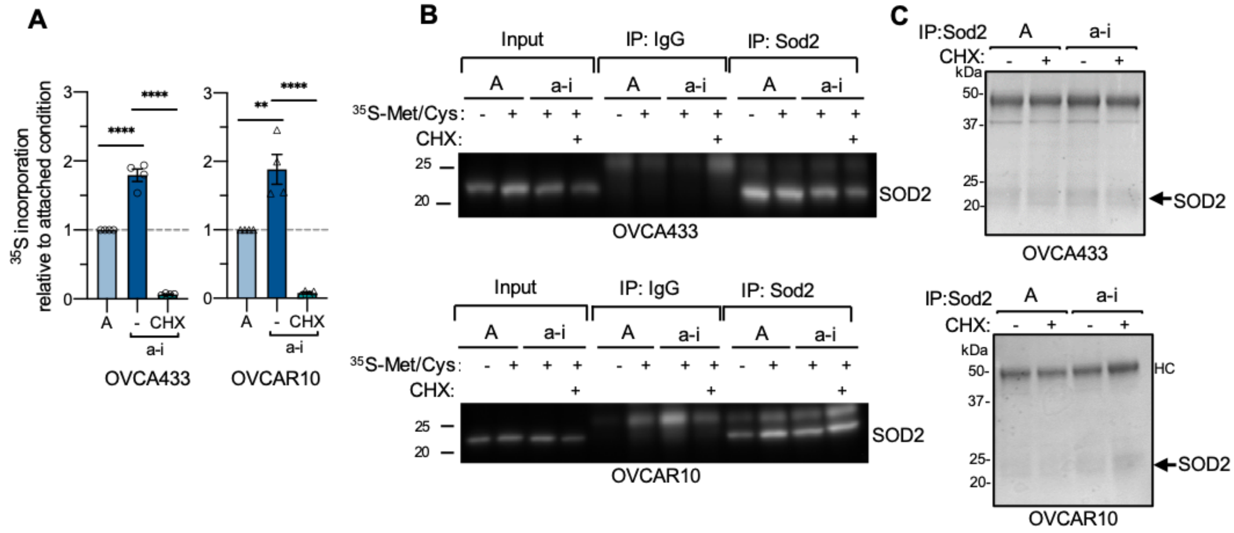


**Supplemental Figure 1.**

A. Global changes in $^{35}$S-Met/Cys incorporation increase under 2 hours in anchorage independent conditions (*n*=4, one-way ANOVA, *P*<0.0001, Tukey's post test ***P*<0.01, *****P*<0.0001).

B. Western blot following IP with control IgG or Sod2 antibodies confirms the specificity of the Sod2 antibody used in C.

C. Following $^{35}$S-Met/Cys incorporation assays in cells cultured in attached (A) or anchorage independent conditions (a-i, 2 h), SOD2 was immunoprecipitated and extracted from SDS-PAGE for liquid scintillation counting (results shown in Fig 1E).

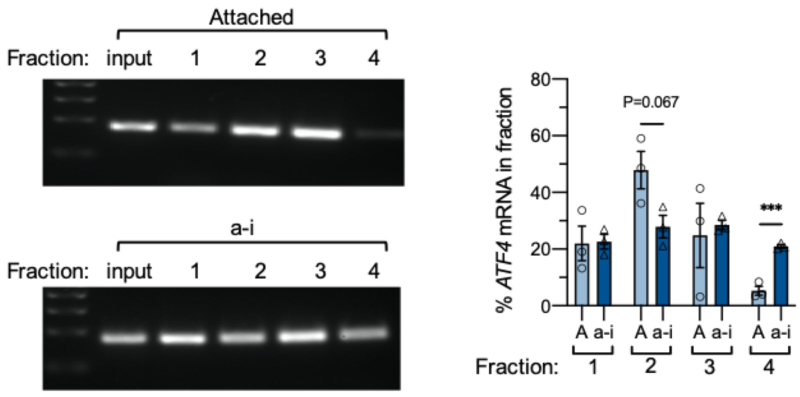


**Supplemental Figure 2.**
Representative RT-PCR and quantification of relative *ATF4* mRNA levels in polysomal fractions isolated from OVCA433 cells cultured in attached (A) or anchorage independent (a-i; 0.5 h) culture conditions (*n*=3; t-test ***$P<0.001$).

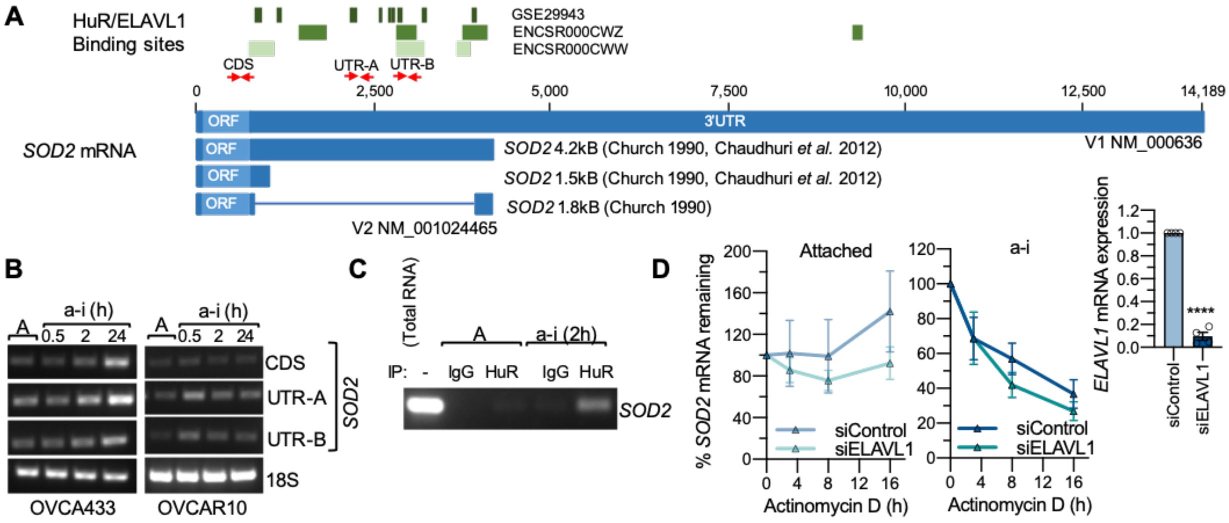


**Supplemental Figure 3.**

A. Several *SOD2* transcripts with alternate 3' UTR splicing and lengths have been reported. Schematic indicates HuR binding sites, as identified by RIP-seq and PAR-CLIP studies.
B. Red arrows in A. indicate primers used to determine the presence of a longer *SOD2* 3'UTR containing HuR binding site in ovarian cancer cell lines under attached (A) and anchorage independent (a-i) conditions.
C. Ribonucleoprotein Immunoprecipitation of HuR and *SOD2* RT-PCR demonstrates that anchorage-independence (a-i, 2h) induces HuR binding to *SOD2* mRNA in OVCAR10 cells
D. HuR knock-down does not affect *SOD2* mRNA stability in attached or anchorage-independent conditions, as determined by Actinomycin D treatment (*n*=4; two-way ANOVA: ns). HuR knock-down was assessed by semi quantitative real time RT-PCR (t-test, ****$P<0.0001$).
